# The first discovery of SFTSV in the Centre of Metropolitan Beijing, China

**DOI:** 10.1101/2024.02.18.580844

**Authors:** Fei Yuan, Lianglong Zhu, Di Tian, Mengyu Xia, Minghao Zheng, Qing Zhang, Tingyu Zhang, Xing Zhang, Aihua Zheng

## Abstract

Severe fever with thrombocytopenia virus (SFTSV), an emerging tick-borne bandavirus, poses a significant public health threat in rural China. Since 2021, an increase in local cases has been noted in the rural-urban fringe surrounding Beijing. This study aimed to assess the formation of natural foci in urban areas by conducting a field survey of ticks and hedgehogs from Beijing’s second to fifth ring roads. Our survey revealed a diverse tick population in city parks, including the major SFTSV vector, *H. longicornis*. Parthenogenetic *H. longicornis*, known for its role in rapid SFTSV spread, was identified in key locations like Beihai Park and Taoranting Park, near the Forbidden City. Notably, high SFTSV seroprevalence and RNA prevalence were found in hedgehogs and parasitic ticks in the center of Beijing. These findings highlight the circulation of SFTSV in central Beijing, underscoring the need for urgent attention and enhanced surveillance measures.

## Introduction

Severe fever with thrombocytopenia syndrome virus (SFTSV) emerged as a novel bandavirus in 2009, initially discovered in the Dabie Mountains at the convergence of Henan, Anhui, and Hubei provinces of China [1]. Subsequently it was reported in South Korea [2], Japan [3], Vietnam [4], Myanmar [5], Pakistan [6], and Thailand [7]. SFTSV infection elicits severe symptoms in patients, encompassing fever, gastrointestinal manifestations, thrombocytopenia, and leukopenia, with reported fatality rates ranging from 6% to 30% across various studies [1,8,9].

Primarily transmitted by the tick species *H. longicornis*, with *H. flava* identified as a potential vector [10], there has been a growing number of reports documenting human-to-human and animal-to-human transmission via close contact [11–15], a phenomenon corroborated by ferret animal models [16]. Notably, *H. longicornis* comprises both bisexual and parthenogenetic populations, with the latter demonstrating accelerated spread and significant responsibility for the rapid spread of SFTSV [17]. Additionally, accumulating evidence underscores the substantial role played by migratory birds in the long-distance dispersal of both *H. longicornis* and SFTSV [17–19]. High seroprevalence rates have been observed across diverse wild and domestic animal species, including sheep (69.5%), cattle (60.4%), dogs (37.9%), chickens (47.4%), as well as shrews, rodents, and hedgehogs [20]. Among the investigated mammalian hosts, hedgehogs emerge as prominent wildlife amplifying hosts capable of sustaining SFTSV transmission via *H. longicornis* bites [21].

Beijing, boasting a history spanning 3,000 years and serving as China’s capital for 800 years since the Jin dynasty in AD 1151, presents a distinctive urban landscape. The cityscape ranges from the iconic Forbidden City at its core to the surrounding periphery encircled by expressways, namely the 2^nd^, 3^rd^, 4^th^, 5^th^, and 6^th^ ring roads. Within the 2^nd^ ring road lies the historic core of the Ming and Qing dynasties, established in AD 1420. Urban Beijing comprises six districts, namely Dongcheng, Xicheng, Haidian, Chaoyang, Fengtai, and Shijingshan, with the majority situated within the 5^th^ ring road. Part of areas between the 4^th^ and 5^th^ ring roads, and areas between the 5^th^ and 6^th^ ring roads, constitute the rural-urban fringe. Additionally, Beijing encompasses ten satellite cities including Shunyi, Mentougou, Pinggu, Tongzhou, Miyun, Huairou, Yanqing, Changping, Fangshan, and Daxing, predominantly located beyond the 5^th^ ring road (Figure S1).

With one of the world’s highest population densities, urban Beijing accommodates approximately 7,943 individuals per square kilometer, reaching densities exceeding 19,000 and 20,000 per square kilometer within the 2^nd^ ring road and between the 2^nd^ and 3^rd^ ring roads, respectively (Beijing 2022 Year Book). Despite its predominantly flat terrain, urban Beijing is encircled by mountainous regions to the west, north, and east, with elevations ranging from 1,000 to 1,500 meters. Positioned along the migration route between East Asia and Australia, Beijing’s avian biodiversity has flourished alongside the expansion of urban green spaces, with a reported 508 bird species documented as of 2023.

The first local human SFTS case in Beijing was reported in October 2021, involving a 69-year-old individual residing in Longquanwu Village, Mentougou District[22,23]. Subsequent natural foci of SFTSV were identified in Longquanwu Village, Mentougou District, and Wanwanshu, Shunyi District, in 2023 [21].

## Materials and Methods

### Ethics statement

All animal studies were carried out in strict accordance with the recommendations in the Guide for the Care and Use of Laboratory Animals of the Ministry of Science and Technology of the People’s Republic of China. The protocols for animal studies were approved by the Committee on the Ethics of Animal Experiments of the Institute of Zoology, Chinese Academy of Sciences (Approval number: IOZ20180058). Human serum samples from SFTS patients were obtained from Beijing Ditan Hospital, Capital Medical University, Beijing, China. A written consent form was obtained from the patients before they participated in this study.

### Field Survey

Hedgehogs were captured in parks and gardens throughout Beijing at night, between July and August 2023. Following anesthesia, approximately 0.5 ml of blood was drawn from the heart of each hedgehog, after which ticks were collected using forceps. Additionally, ticks in grassy areas were also collected using flag-dragging method at several locations. All ticks were taxonomically identified under a dissecting microscope based on their morphological characteristics. Blood was separated into serum and blood cells and kept at -80°C.

### Phylogenetic analysis of ticks

One leg from each tick were removed for analysis. Parthenogenetic ticks were determined by the deletion of a single base T at nucleotide position 8497 in the untranslated region of the mitochondrial genome[24]. Phylogenetic analysis was performed using the full-length mitochondrial genomes. Tick DNA was extracted using the MightyPrep reagent for DNA Kit (Takara, Japan) according to the manufacturer’s instructions. The mitochondrial DNA were sequenced by next generation sequencing (Tsingke Biotech, Beijing, China) and deposited in GenBank. Mitochondrial genomes of *H. longicornis* ticks from SFTS endemic areas were included in Table S1.

Maximum likelihood tree was constructed by using the maximum likelihood method, MEGA-X with the bootstrap value set at 1000. For Bayesian-inference (BI) method, Best-fit models for Bayesian-inference (BI) method were selected by MrModeltest ver. 2.3; GTR + I + G model was chosen for all datasets according to the Akaike Information Criterion (AIC). the Markov chain Monte Carlo (MCMC) algorithm was run up to 1,000, 000 generations. Samples were taken every 1,000 generations, with a burn-in set to 25%. The average split frequencies between the runs were less than 1%. The two final trees were viewed and edited by FigTree ver. 1.4. 3 and Photoshop CC 2018.

### RNA extraction and amplification of SFTSV RNA by Nested PCR

Total RNAs were extracted from tick homogenates and hedgehog blood cells utilizing either TRIzol reagent (Thermo Fisher Scientific, USA) or the RNeasy kit (Qiagen, Germany) in accordance with the respective manufacturer’s protocols. The RNA samples were then reverse transcribed employing a One-Step SYBR PrimerScript reverse transcription (RT)-PCR kit (TaKaRa, Japan). Nested PCR conditions were as follows: initial denaturation at 95°C for 3 min, followed by denaturation at 95°C for 10 sec, 40 cycles of amplification at 95°C for 5 sec, and annealing/extension at 60°C for 30 sec. Primer sequences utilized are detailed in Table S2. The PCR products were sequenced by Sanger method and deposited in GenBase (Accession number C_AA059120.1-C_AA059155.1).

SFTSV sequences retrieved from prior studies were acquired from GenBank (1,2) (Table S3). Phylogenetic trees were constructed utilizing the maximum likelihood method within the MEGA program. The robustness of the resulting tree topologies was assessed via 1000 bootstrap replications to ascertain confidence levels.

### Neutralizing assay

The neutralization titers of the sera samples obtained from hedgehogs and patients was evaluated using Focus Reduction Neutralization Test (FRNT). In brief, a fixed amount of rVSV-GFP-SFTSV AH12 strain (∼1000 FFU) was mixed with heat-inactivated sera samples, which were serially diluted five-fold from 1:20 to 1:2500. After a 30-minute incubation at room temperature, the virus-serum mixtures were added onto Vero cells in 96-well plates. The plates were then incubated at 37°C for two hours to allow virus entry and infection. Subsequently, the supernatants were replaced with fresh DMEM containing 2% FBS and 20 mM NH_4_Cl, and the plates were further incubated at 28°C for 24 hours. The number of GFP-positive cells was counted and neutralization titers were calculated using the Reed-Muench method. The cut-off value of 30 was determined by naïve Hedgehog samples.

## Results

### Case reports of three local SFTS patients

In 2023, six local cases of SFTS were reported in Beijing, with four residing in Beishiqu, Houjiazhuang, and Donglujiao Village in Pinggu district, one in Baixi Village in Fengtai District, and one in Gaoying Village in Tongzhou District (China CDC) (Figure S1). Among these, three patients aged 50 to 61 from Beishiqu, Houjiazhuang, and Gaoying were admitted to Beijing Ditan Hospital. All three patients tested positive for SFTSV by Realtime PCR, with two having a history of tick bites. Common symptoms included fever, altered consciousness, and slurred speech, accompanied by leukopenia and thrombocytopenia. Additionally, varying degrees of organ failure, including encephalitis, cardiac damage, liver injury, renal impairment, and increased pancreatic enzymes, were observed. Treatment with favipiravir antiviral medication, dexamethasone, and supportive care for two weeks led to the recovery and subsequent discharge of all patients. Detailed blood test results are depicted in Figure 1.

**Figure 1.**
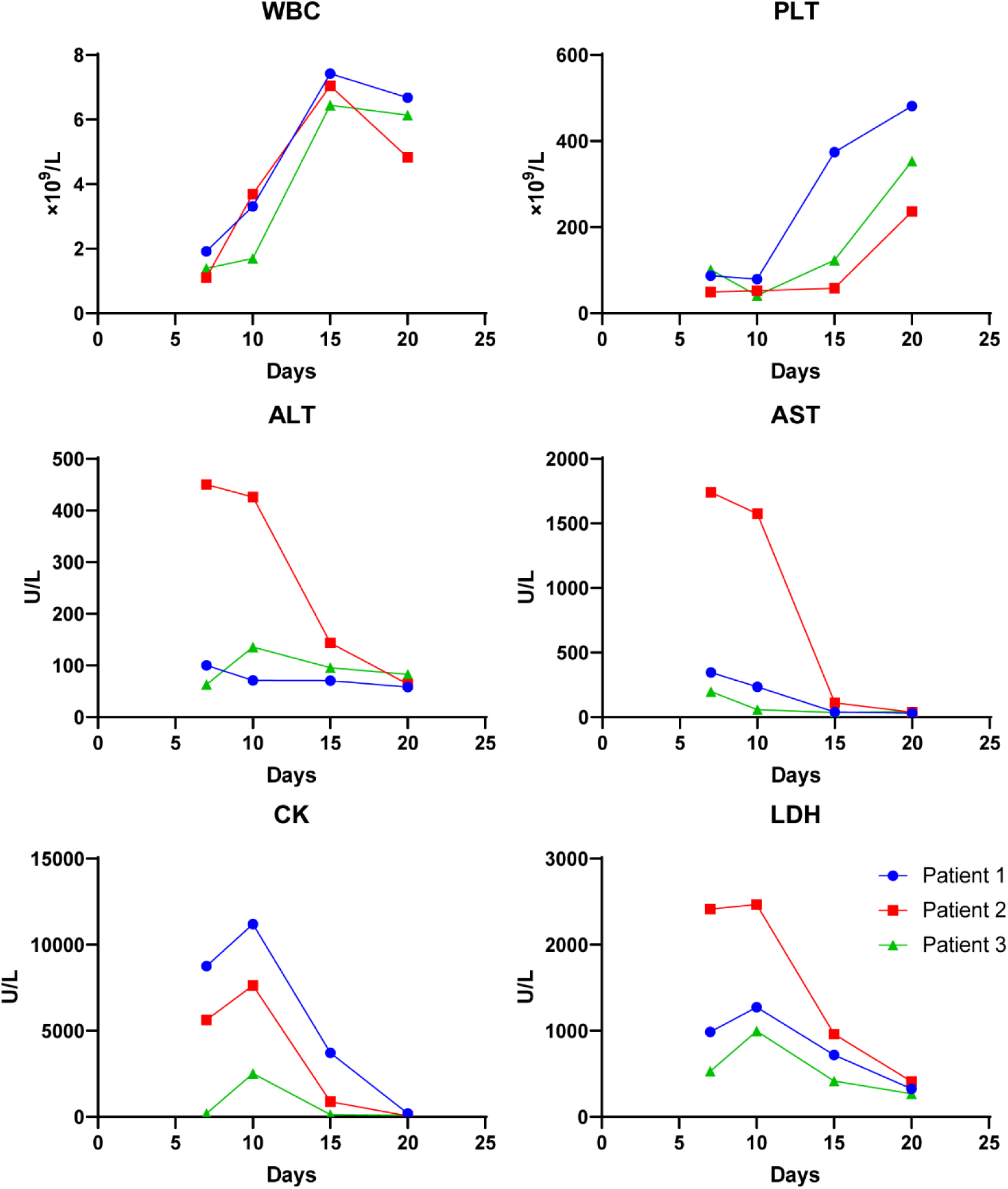
Blood test results of three patients. White blood cell count (WBC), platelet count (PLT), alanine aminotransferase (ALT), aspartate aminotransferase (AST), creatine kinase (CK), and lactate dehydrogenase (LDH) levels are depicted for each patient over the course of days following symptom onset.

Notably, a near-complete-length genome was identified from the serum of the Gaoying patient, while only short segments were obtained from the sera of the other two patients.

### Diversity of ticks in Beijing

All reported human SFTS cases thus far have been localized in the rural-urban fringe areas surrounding Beijing (Figure S1). To assess the potential establishment of natural SFTSV foci in urban areas, a field survey was conducted from July to August 2023. Host-seeking ticks were collected from vegetation at 6 sites, and parasitic ticks were collected from hedgehogs at 21 sites, covering patient residences and major parks or universities within the 5^th^ ring road (Figure 2 and Table 1). Analysis revealed that in the rural-urban fringe areas outside the 5^th^ ring road, *H. longicornis* was the dominant tick species, followed by *H. flava*, *Rhipicephalus sanguineus*, and *Dermacentor* spp. Parthenogenetic *H. longicornis* were detected in Beijing Wildlife Rescue and Rehabilitation Center, Gaoying Village, and Beishiqu Village, with the latter two locations in close proximity to patient residences. Contrary to previous assumptions, urban areas, particularly within the 4^th^ ring road, exhibited high diversity and abundance of ticks, predominantly *R. sanguineus*, followed by *H. flava*, with few *Dermacentor* spp. and *H. longicornis* (Figure 2). The total tick infestation on hedgehogs reached as high as 136 per animal, with no significant difference observed from the 2^nd^ ring to the 5^th^ ring road (Table 2). *H. longicornis* ticks were detected in Laoshan Park, Olympic Forest Park, Wanliu, Beihai Park, and Taoranting Park, with numbers of 4, 18, 22, 10, and 2 respectively. Parthenogenetic *H. longicornis* was identified in Olympic Forest Park, Wanliu, Beihai Park, and Taoranting Park, the latter two located inside the 2^nd^ ring road adjacent to the Forbidden City (Figure 2). Phylogenetic analysis of whole mitochondrial sequences of the parthenogenetic ticks revealed three distinct clusters, namely Beishiqu, Wanwanshu/Olympic Forest Park/Beijing Wildlife Rescue and Rehabilitation Center, and Gaoying/Beihai/Taoranting, indicating the invasion of at least three strains of parthenogenetic *H. longicornis* into Beijing.

**Figure 2.**
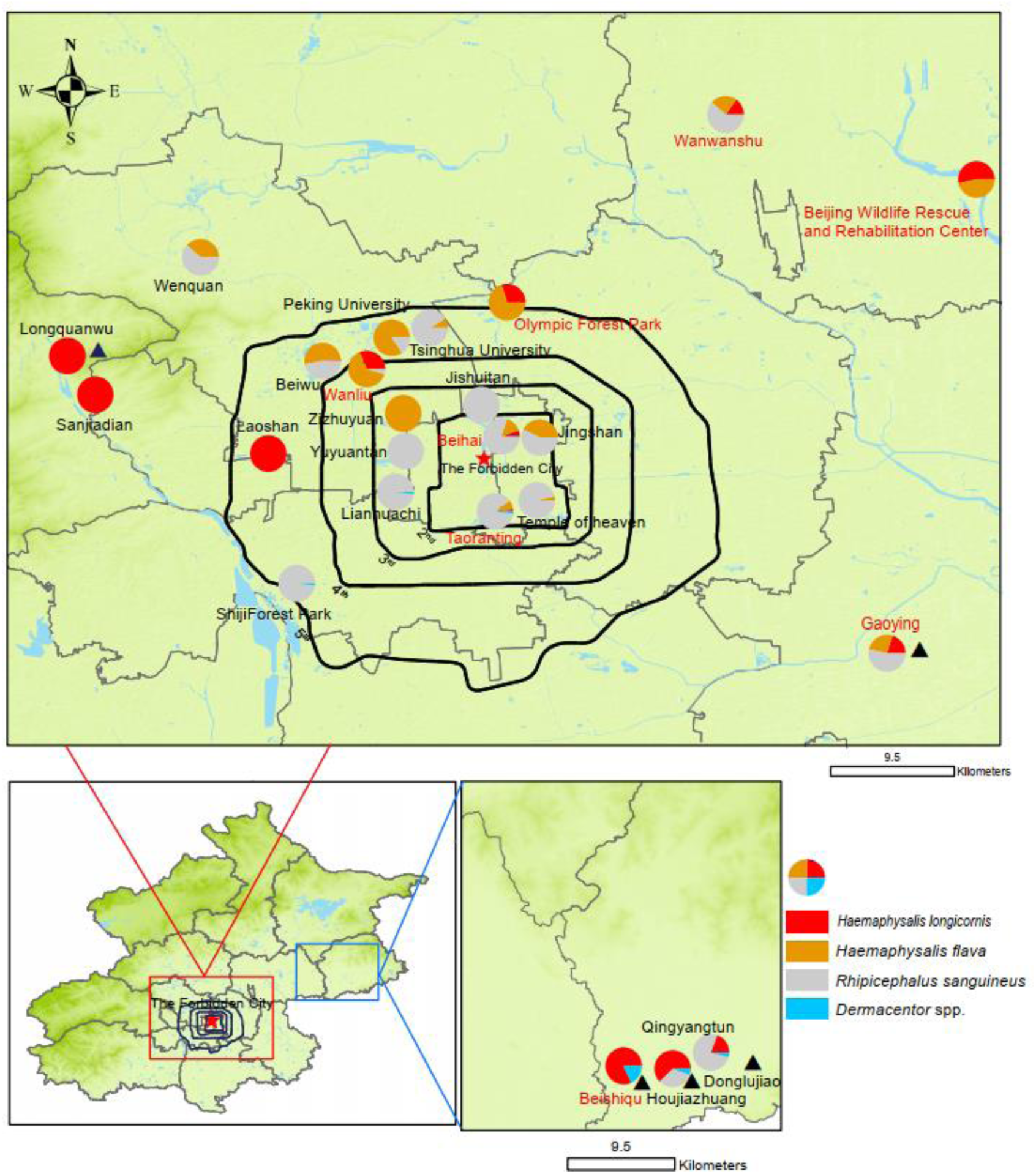
Tick diversity in Beijing. The map was created by ArcMap 10.8 (ArcGIS Enterprise, ESRI, Redlands, CA, USA). The pie chart illustrates tick species distribution, with red font denoting sampling points containing parthenogenetic *H. longicornis*, and black triangles indicating locations of confirmed SFTS cases. Ring roads are delineated in black.

**Figure 3.**
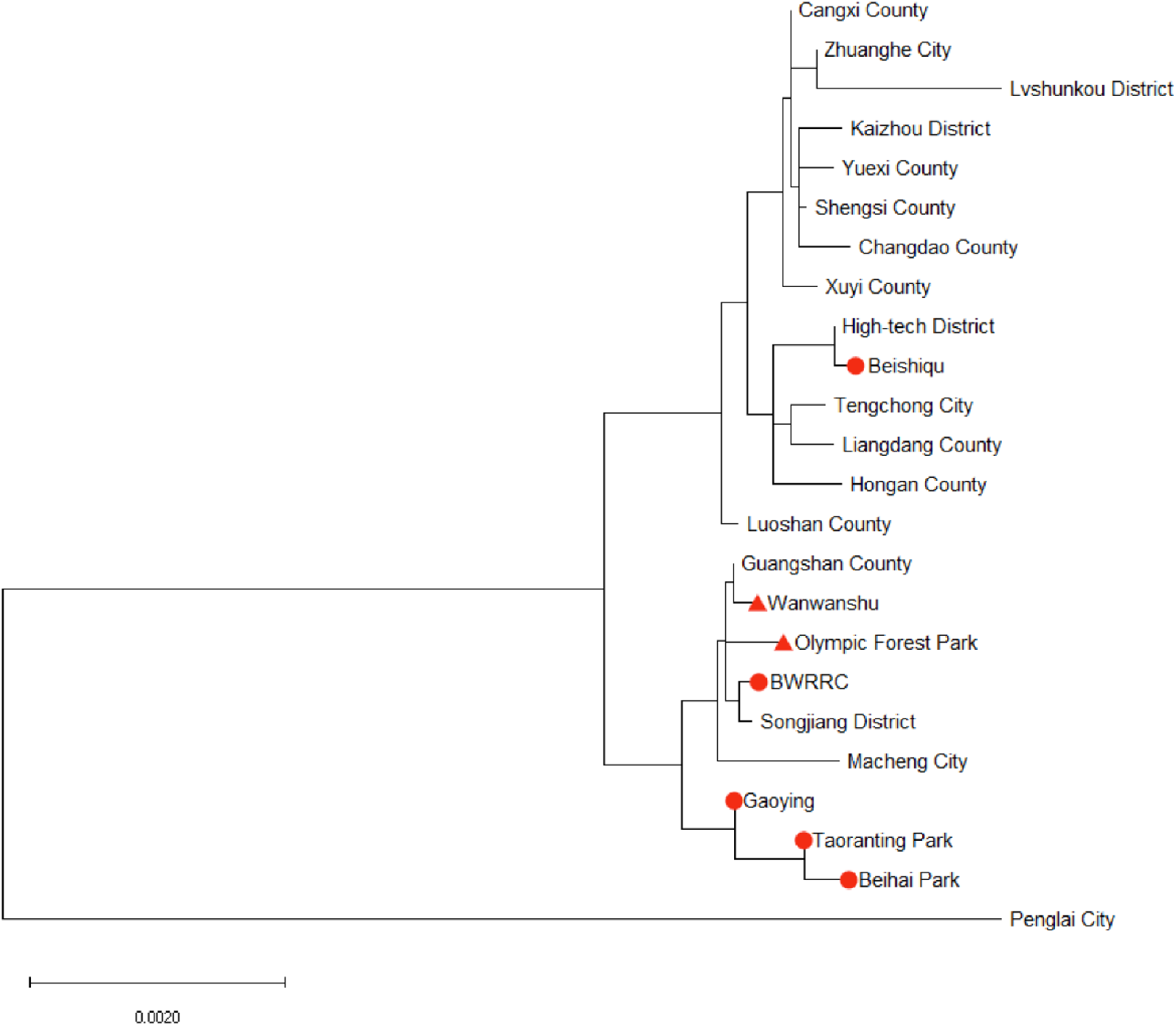
Phylogenetic analysis of the *H. longicornis* parthenogenetic population. Maximum likelihood tree was established with the mitochondrial genomes of *H. longicornis* collected in Beijing (red circle collected in 2023, red triangle collected in 2021) and from SFTS endemic areas. Constructed by MEGA-X with the bootstrap value set at 1000.

**Table 1.** Epidemiological analysis of ticks and hedgehogs. Locations were categorized by the 2^nd^ to 5^th^ ring road of Beijing City. Hedgehog SFTSV RNA Positive Rate (Number of positive by PCR/number tested); Hedgehog SFTSV antibody Positive Rate (Number of positive by neutralizing assay/number tested); Hedgehog infested tick RNA SFTSV RNA Positive Rate was tested by pooling the ticks from individual hedgehog and then detecting SFTSV RNA by PCR (Number of positive by PCR/number tested); SFTSV RNA in tick on Vegetation, tick pools were tested by PCR. +, positive; -, negative. NA, not available. All the PCR products of SFTSV RNA was sequenced by Sanger sequencing method. No., number. BWRRC, Beijing Wildlife Rescue and Rehabilitation Center. Univ., university.

| Location | Hedgehog<br>SFTSV RNA<br>Positive Rate | Hedgehog<br>SFTSV antibody<br>Positive Rate | Tick no. per<br>hedgehog<br>(average) | Hedgehog infested<br>tick RNA SFTSV<br>RNA Positive Rate | SFTSV<br>RNA in tick on<br>Vegetation |
| --- | --- | --- | --- | --- | --- |
| **<2 °ring** |  |  |  |  |  |
| Jingshan | 1/2 | 1/2 | 50 | 1/2 | NA |
| Beihai | 3/4 | 3/4 | 108 | 1/4 | NA |
| Taoranting | NA | 1/4 | 76 | NA | NA |
| Temple of heaven | 2/5 | 4/5 | 130 | 0/5 | NA |
| **2 °- 3 °ring** |  |  |  |  |  |
| Zizhuyuan | 2/2 | 2/5 | 40 | 1/2 | NA |
| Jishuitan | 2/2 | 1/2 | 43 | NA | NA |
| Lianhuachi | 1/2 | 0/1 | 102 | NA | NA |
| Yuyuantan | 0/1 | 0/1 | 20 | 0/1 | NA |
| **3 °- 4 °ring** |  |  |  |  |  |
| Wanliu | 3/4 | 1/4 | 25 | 1/4 | NA |
| **4 °- 5 °ring** |  |  |  |  |  |
| Laoshan | NA | NA | 25 | NA | - |
| Olympic Forest Park | 0/1 | 2/2 | 105 | 0/1 | - |
| Peking Univ. | 0/1 | NA | 26 | 0/1 | NA |
| Tsinghua Univ. | 0/2 | 0/1 | 23 | 0/2 | NA |
| Beiwu | 0/2 | 0/2 | 32 | 1/2 | + |
| Shiji Forest Park | 1/2 | 2/2 | 112 | NA | NA |
| **>5 °ring** |  |  |  |  |  |
| Beishiqu | NA | 1/1 | 63 | NA | NA |
| Houjiazhuang | 2/4 | 2/2 | 74 | 4/4 | NA |
| Qingyangtun | 0/2 | 1/2 | 66 | 1/2 | NA |
| Wenquan | 1/3 | 1/3 | 52 | NA | NA |
| Gaoying | 0/2 | 2/3 | 4 | 0/2 | + |
| BWRRC | NA | NA | NA | NA | + |

**Table 2.** Density of hedgehogs and number of stray cats in the parks. The density of hedgehogs was calculated by number of captured animals divided by the area searched from 7 to 9 pm (no. of animals per km^2^). There are two limitations in this method: 1) hedgehogs are more active after 11 pm during the night; 2) We only searched the bush and woods area. The number of stray cats was obtained by interviewing the volunteers feeding the cats. †Urban; ‡ rural–urban fringe area. NA, not available.

| Location | Hedgehog density | Cat no. |
| --- | --- | --- |
| Jingshan† | NA | 9 |
| Beihai† | 360 | 15 |
| Taoranting† | 500 | 10 |
| Temple of heaven† | 130 | 20 |
| Zizhuyuan† | 200 | NA |
| Lianhuachi† | NA | 5 |
| Yuyuantan† | NA | 15 |
| Wanliu† | 300 | NA |
| Beishiqu‡ | 200 | NA |
| Houjiazhuang‡ | 600 | NA |
| Qingyangtun‡ | 400 | NA |
| Wenquan‡ | 300 | NA |
| Longquanwu‡ | 250 | NA |

### SFTSV prevalence in ticks and hedgehogs

Fifty hedgehogs from 18 locations spanning the 2^nd^ to 5^th^ ring road and patient residences were assayed for SFTSV seroprevalence against recombinant SFTSV AH12 strain. Twenty-eight hedgehogs from 15 locations tested positive for SFTSV neutralizing antibodies, with the highest titer of 1000 detected at Houjiazhuang Village in Pinggu District, adjacent to a patient’s residence. SFTSV-neutralizing antibodies were also detected in hedgehogs within the 5^th^ ring road, ranging in titers from the cutoff of 30 to 250. Notably, positive hedgehogs were detected in all four locations within the 2^nd^ ring road (Table 1 and Figure 4). Subsequently, SFTSV RNA was tested in blood samples from hedgehogs, host-seeking ticks, and hedgehog-parasitic ticks. SFTSV RNA was amplified from 19 of 46 blood samples tested, with three positive locations identified within the 2^nd^ ring road. Among the pooled parasitic tick samples, 11 out of 37 pools tested positive for SFTSV RNA. SFTSV RNA was detected across the 2^nd^ to 5^th^ ring road of urban Beijing, with three out of five pools of host-seeking ticks testing positive (Table 1).

**Figure 4.**
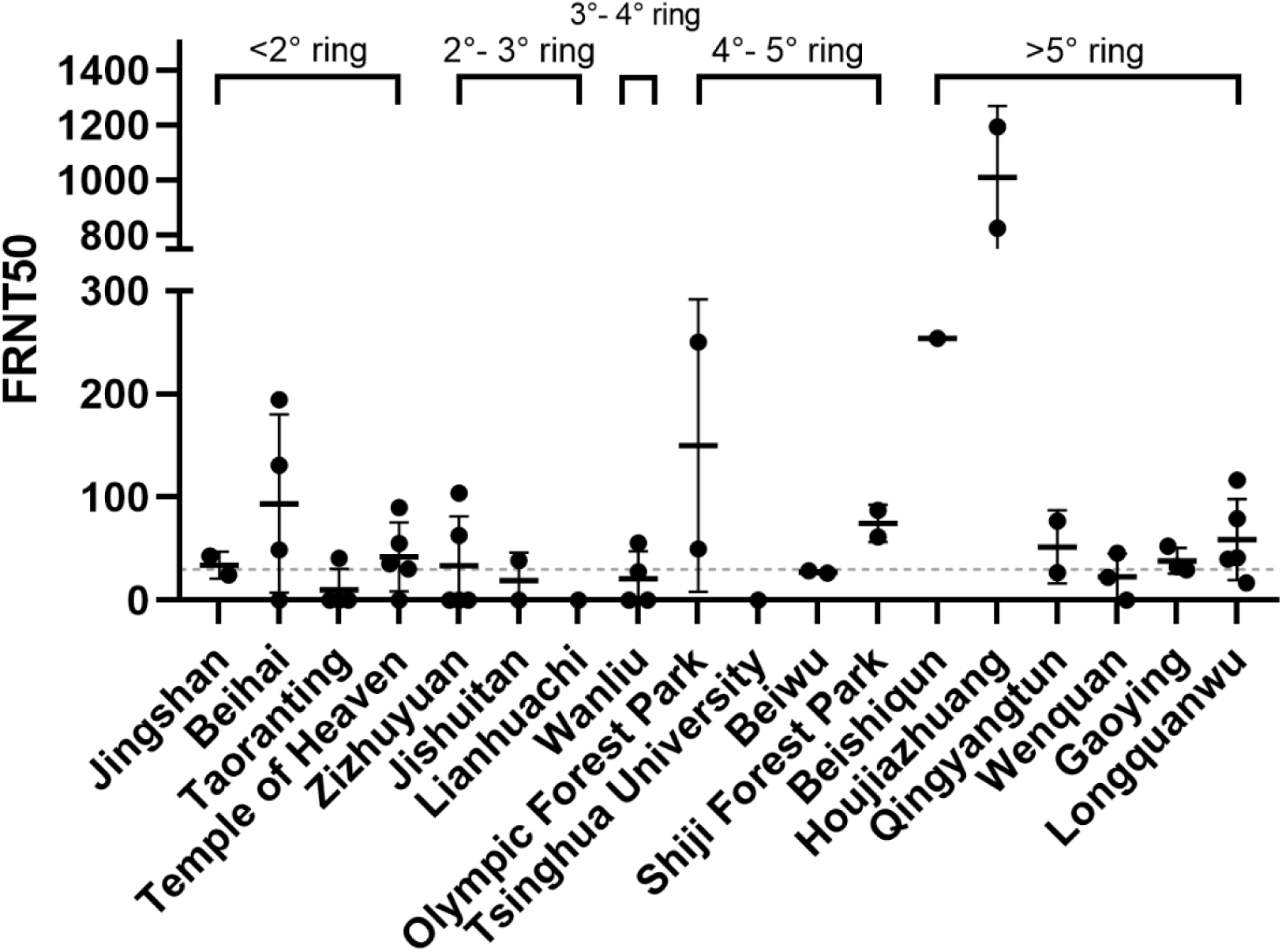
Neutralizing antibody titers (FRNT50) in hedgehogs from various. Locations in Beijing Neutralizing antibody titers against rVSV-GFP-SFTSV AH12 strain in hedgehogs captured at specified locations in Beijing are presented. The dashed line indicates the cutoff value set at 30.

All amplified SFTSV RNA samples were sequenced using the Sanger method. Two near full-length genomes were identified from parasitic ticks in Houjiazhuang Village and Beishiqu Village in Pinggu District, near patient residences. However, only partial sequences of S (∼200bp), M and L segments (∼800 bp) were obtained from samples collected within the 5^th^ ring road. Phylogenetic analysis revealed that the three near-complete length SFTSV strains belonged to the C2 lineage, as aligned with partial sequences of M and L segments (Figure 5). Furthermore, alignment with partial sequences of M or L segments indicated that all SFTSV strains within the 5^th^ ring road belonged to the C2 lineage (Figure 5A and 5C), with a single strain from Wenquan Village in Haidian District, located in the rural-urban fringe area, falling into the C4 lineage (Figure 5B and Figure 2). These findings suggest the circulation of multiple SFTSV lineages in Beijing City.

**Figure 5.**
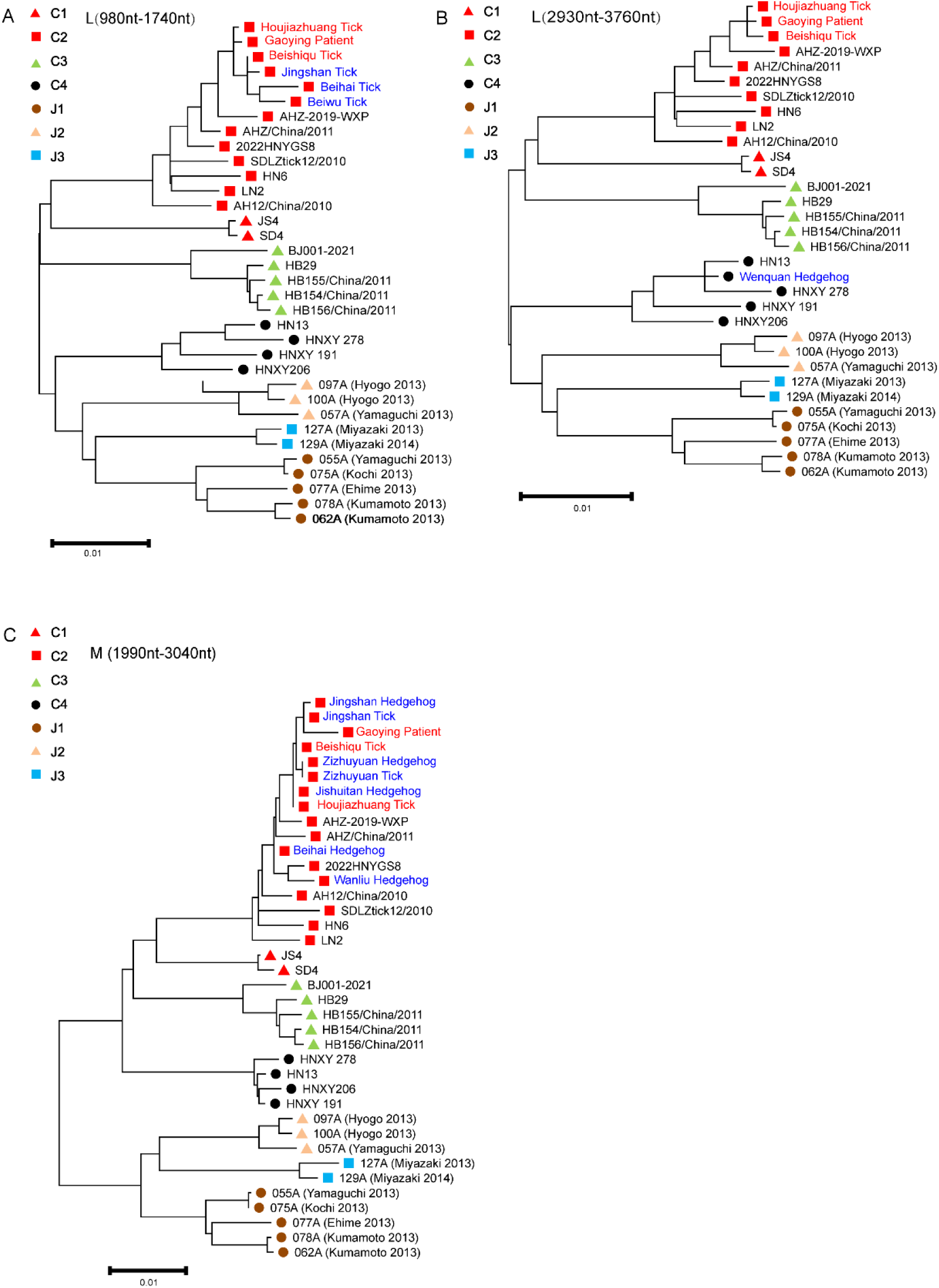
Phylogenetic analysis of SFTSV isolates. A maximum likelihood tree was constructed based on partial L or M segments of SFTSV isolates (Table S3). SFTSV lineages are represented by colors and shapes. Red font signifies near-complete SFTSV genome sequences, while blue font indicates partial sequences of the SFTSV genome.

## Discussion

Our previous research on the vectors and hosts of SFTSV yielded two hypotheses. Firstly, parthenogenetic *H. longicornis* ticks are strongly associated with SFTSV outbreaks and serve as early indicators of SFTSV emergence in new areas [17]. Secondly, SFTSV transmission can be sustained between *H. longicornis* ticks and hedgehogs in urban areas across China [21,25].

Despite the rapid spread of SFTSV across China and Asia, no local cases were reported in Beijing following its initial discovery in 2009. However, in 2019, parthenogenetic *H. longicornis* ticks were first identified in two rural-urban locations: Wanwanshu in the Shunyi district and Olympic Forest Park in the Chaoyang District [17]. Subsequently, SFTSV RNA was detected in *H. longicornis* collected from Wanwanshu in spring 2021. In fall 2021, the first SFTS patient was diagnosed in Longquanwu village, Mentougou District, located 26 miles away from Wanwanshu. In 2023, six human cases had been reported, with four residing in close proximity within three villages in Pinggu District, one in Gaoying village, Tongzhou District, and one in Baixi Village Fengtai District. Our field survey confirmed the presence of parthenogenetic *H. longicornis* ticks in Beshiqu and Gaoying Village, further supporting their association with SFTSV transmission. Additionally, parthenogenetic ticks were detected within the 4^th^ ring road in Wanliu, Beihai Park, and Taoranting Park, albeit in low abundance, suggesting that urban SFTSV circulation may still be in its early stages.

Hedgehogs emerge as promising sentinel animals for SFTSV surveillance due to their high susceptibility to infection and their propensity to harbor ticks, including *H. longicornis and H. flava*, at densities of up to 130 ticks per animal [25]. Our survey revealed a high ratio of SFTSV RNA and seroprevalence in hedgehogs, as well as SFTSV RNA prevalence in ticks infesting hedgehogs, underscoring their potential role in SFTSV maintenance. Effective control measures targeting *H. longicornis* and hedgehogs are therefore crucial for SFTSV control in urban areas. Further, hedgehogs are easy to capture due to the slow movement.

Beijing is high urbanized, with population density similar as Manhaton, particularly within the 4^th^ ring road. However, the strict ecological conservation efforts and increased greening rates is creating conducive environments for tick infestation. Besides the common tick species *R. sanguineus*, our survey detected *H. longicornis, H. flava, and Dermacentor* spp. The average tick infestation rate on hedgehogs exceeded 100 ticks per animal in 33% of parks surveyed, indicating a high tick density in urban parks. The most common mammals are Amur hedgehogs and Yellow weasels (*Mustela sibirica*). However, the tick-infestation rate and SFTSV seroprevalence of Yellow weasels is low [25]. Stray cats are also present in parks and communities, albeit at lower densities compared to hedgehogs (Table 2). The high ratio of SFTSV RNA and seroprevalence in hedgehogs, coupled with SFTSV RNA prevalence in ticks infesting hedgehogs, suggests potential SFTSV maintenance by *H. longicornis* and hedgehogs. Moreover, the high density and diversity of birds in Beijing, including the Spotted Dove (*Spilopelia chinensis*), known to exhibit high SFTSV viremia upon infection, may facilitate local SFTSV spread and tick dispersal [26].

We propose that parthenogenetic *H. longicornis* and SFTSV were introduced into Beijing via migratory birds, given Beijing’s location on the East Asian-Australasian Flyway and the presence of over 500 bird species. [21]. Phylogenetic analysis of parthenogenetic tick mitochondrial genomes revealed close genetic relationships between ticks from Beijing Wildlife Rescue and Rehabilitation Center, Wanwanshu, and Olympic Forest Park, detected in 2019 [17]. However, strains from Beishiqu, Pinggu District and Gaoying, Tongzhou District, represent two new lineages. The proximity of the Beihai and Taoranting strains within the 2^nd^ ring road to the Gaoying strain near the 6^th^ ring road suggests potential tick spread from the city’s periphery to its center. Phylogenetic analysis of SFTSV RNA indicated the circulation of both C2 and C4 lineages in Beijing, suggesting multiple SFTSV introduction events.

Notably, no SFTS patients have been reported within the 4^th^ ring road thus far. This could be attributed to the low abundance of *H. longicornis* in the city and the amplification of near-complete SFTSV genomes from ticks near patients’ residences in the rural-urban fringe area, as opposed to only partial sequences obtained within urban areas. Additionally, the implementation of the “Keep off the grass” policy has significantly reduced tick exposure among city residents. However, ongoing monitoring of SFTSV seroprevalence among gardeners is warranted. Our observations also revealed hedgehogs feeding on food provided for stray cats, raising concerns regarding potential cat-human transmission of SFTSV, as reported in Japan and China. Therefore, caution regarding contact with cats is advised.

In conclusion, our findings indicate the establishment of urban SFTSV circulation in Beijing, warranting urgent attention and comprehensive control measures.

## Acknowledgments

We gratefully thank the following funders: National Key R&D Program of China (2021YFC2300903 and 2022YFC2601603), and the Open Research Fund Program of State Key Laboratory of Integrated Pest Management (IPM2315). We also thank Mingxi Zheng from Zhongguancun No.1 Primary School for helping collecting hedgehog samples.

---

**Figure S1.**
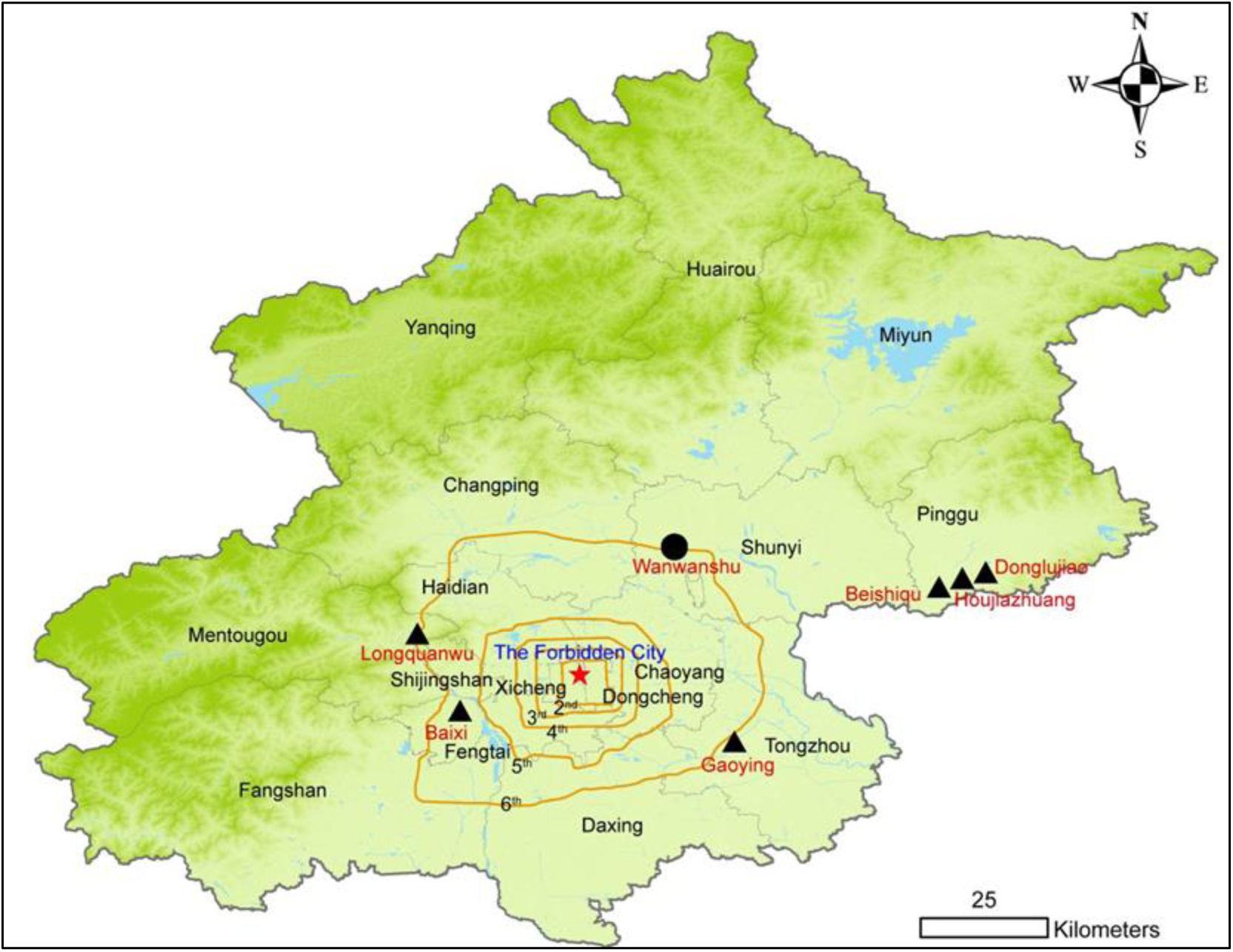
Map of Beijing. Counties and Districts are labeled in black. Red star indicates the center of Beijing, the Forbidden City. Black circle indicates the natural foci of SFTSV. Black triangles indicate the reported patient’s residences and the locations are labeled in red. The 2^nd^ to 6^th^ ring roads are labeled in orange.

**Figure S2.**
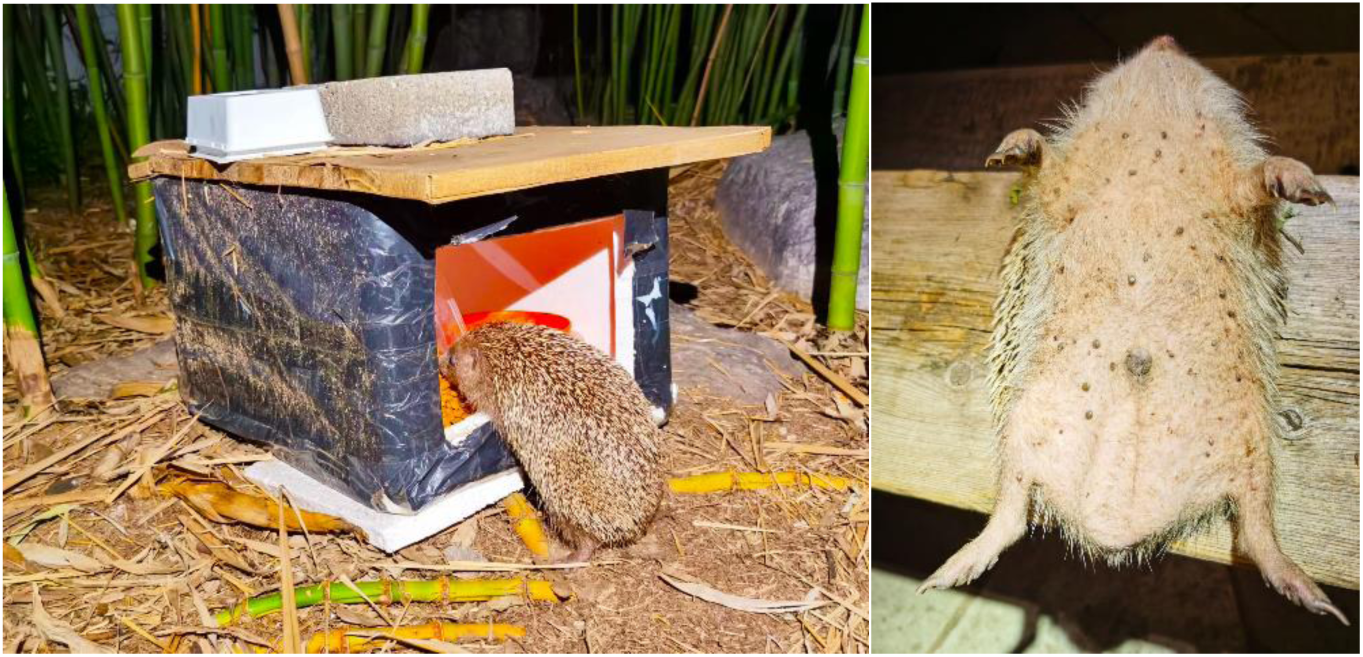
Amur hedgehogs (*Erinaceus amurensis*) in the Zizhuyuan Park. Left, a hedgehog was feeding on the food for stray cats provided by the volunteers. Right, the ticks infested on the belly of an anaesthetized hedgehog.

**Table S1.** Detail information of the mitochondrial genomes of *H. longicornis* ticks.

| Name | Location | Accession number | Year |
| --- | --- | --- | --- |
| Shengsi County | Shengsi County, Zhoushan City, Zhejiang Province | MW642397.1 | 2019 |
| Yuexi County | Yuexi County, Anqing City, Anhui Province | MW642389.1 | 2019 |
| Kaizhou District | Kaizhou District, Chongqing | MW642350.1 | 2019 |
| Changdao County | Penglai District, Yantai City, Shandong Province | MW642340.1 | 2019 |
| Penglai City | Penglai District, Yantai City, Shandong Province | MW642362.1 | 2019 |
| Guidong County | Guidong County, Chenzhou City, Hunan Province | MW642347.1 | 2019 |
| Macheng City | Macheng City, Hubei Province | MW642359.1 | 2019 |
| Cangxi County | Cangxi County, Guangyuan City, Sichuan Province | MW642339.1 | 2019 |
| Xuyi County | Xuyi County, Huai'an City, Jiangsu Province | MW642368.1 | 2019 |
| Lvshunkou District | Lvshunkou District, Dalian City, Liaoning Province | MW642402.1 | 2019 |
| Zhuanghe City | Zhuanghe City, Dalian City, Liaoning Province | MW642399.1 | 2019 |
| High-tech District | High-tech District, Weihai City, Shandong Province | MW642372.1 | 2019 |
| Hanshan County | Hanshan County, Maanshan City, Anhui Province | MW642348.1 | 2019 |
| Hongan County | Hongan County, Huanggang City, Hubei Province | MW642403.1 | 2019 |
| Liangdang County | Liangdang County, Longnan City, Gansu Province | MW642369.1 | 2019 |
| Songjiang District | Songjiang District, Shanghai City | MW642363.1 | 2019 |
| Suichuan County | Suichuan County, Ji 'an City, Jiangxi Province | MW642385.1 | 2019 |
| Tengchong City | Tengchong City, Baoshan City, Yunnan Province | MW642390.1 | 2019 |
| Luoshan County | Luoshan County, Xinyang City, Henan Province | MW642407.1 | 2019 |
| Guangshan County | Guangshan County, Xinyang City, Henan Province | MW642376.1 | 2019 |

**Table S2.**
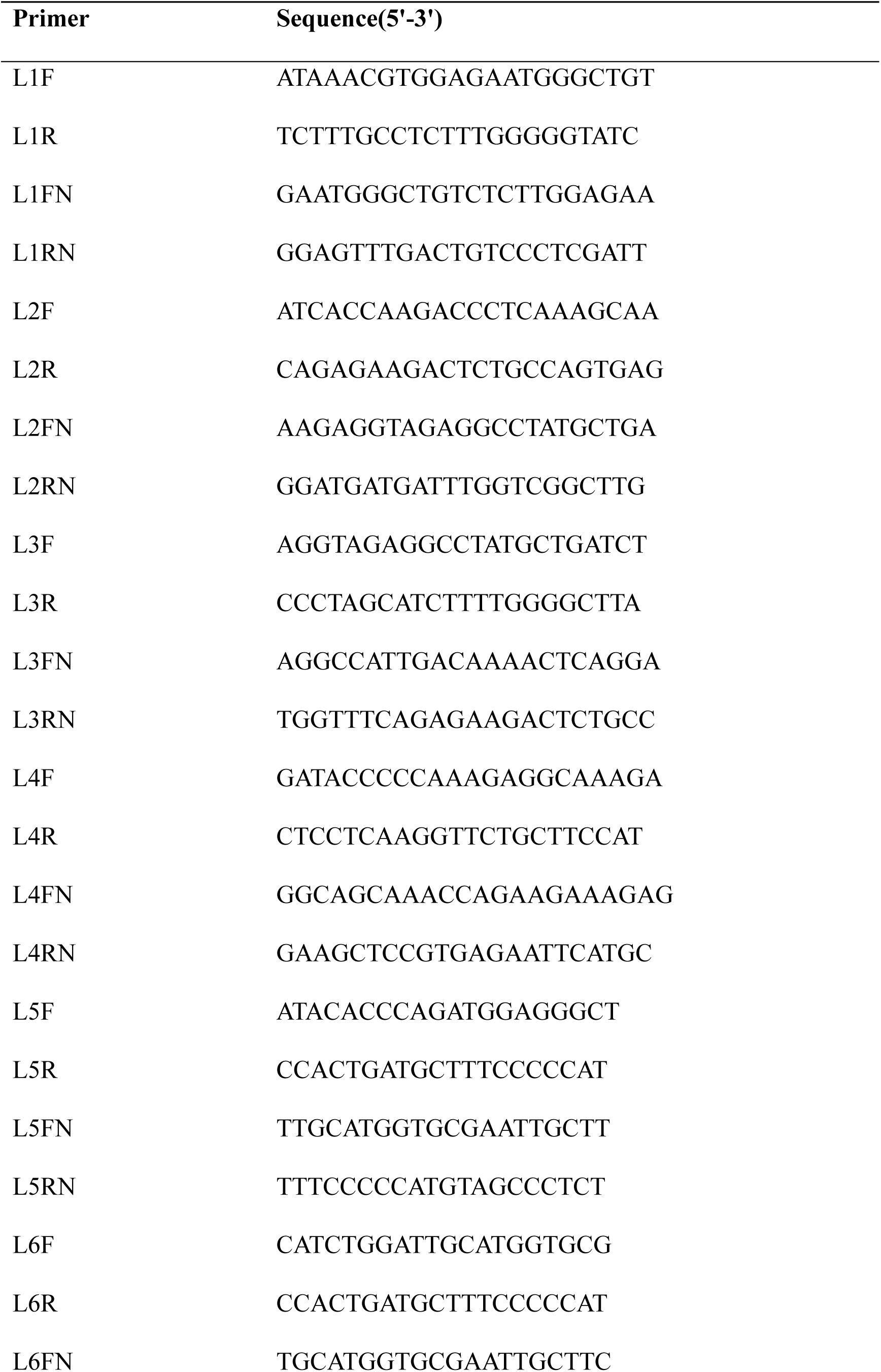
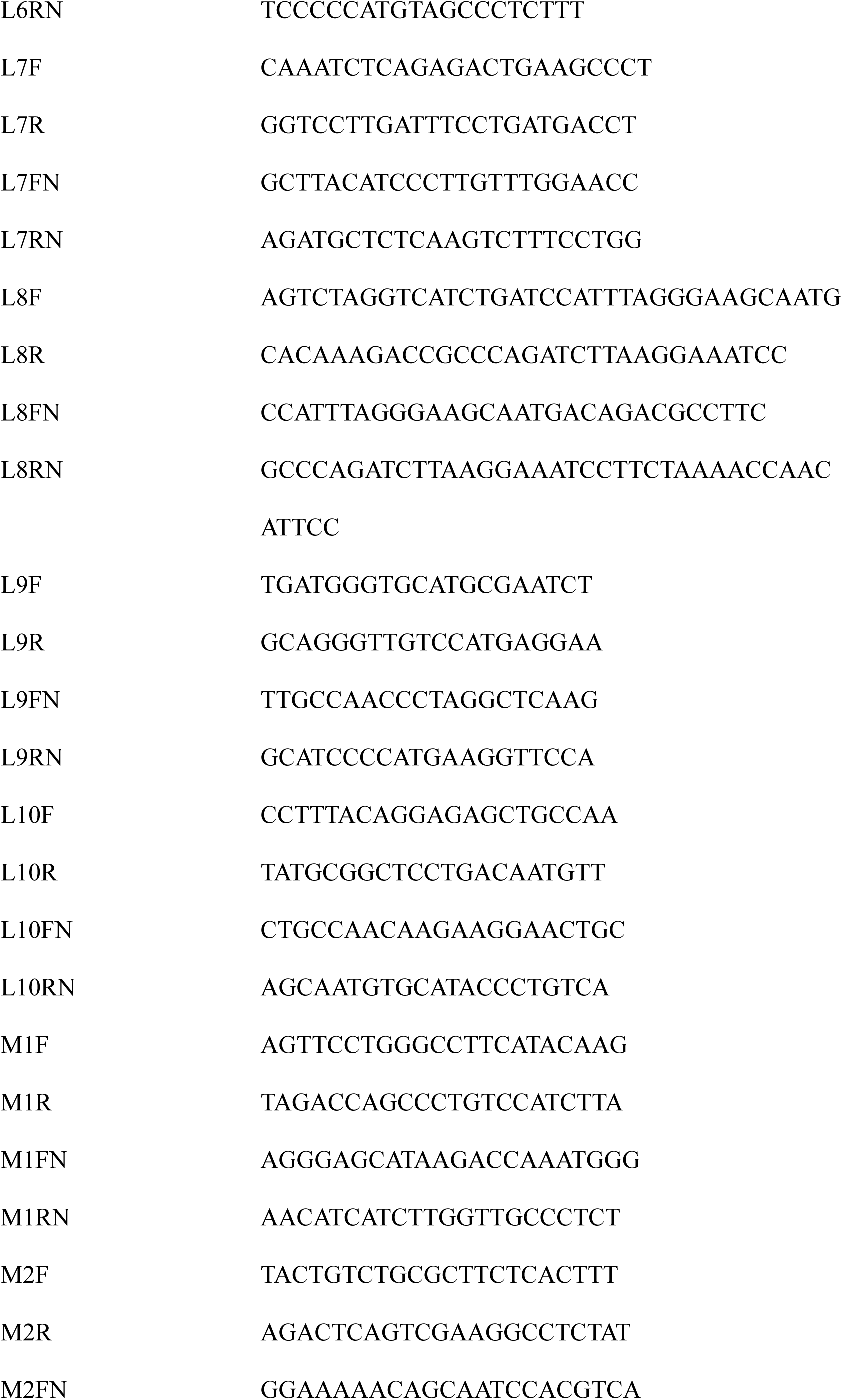
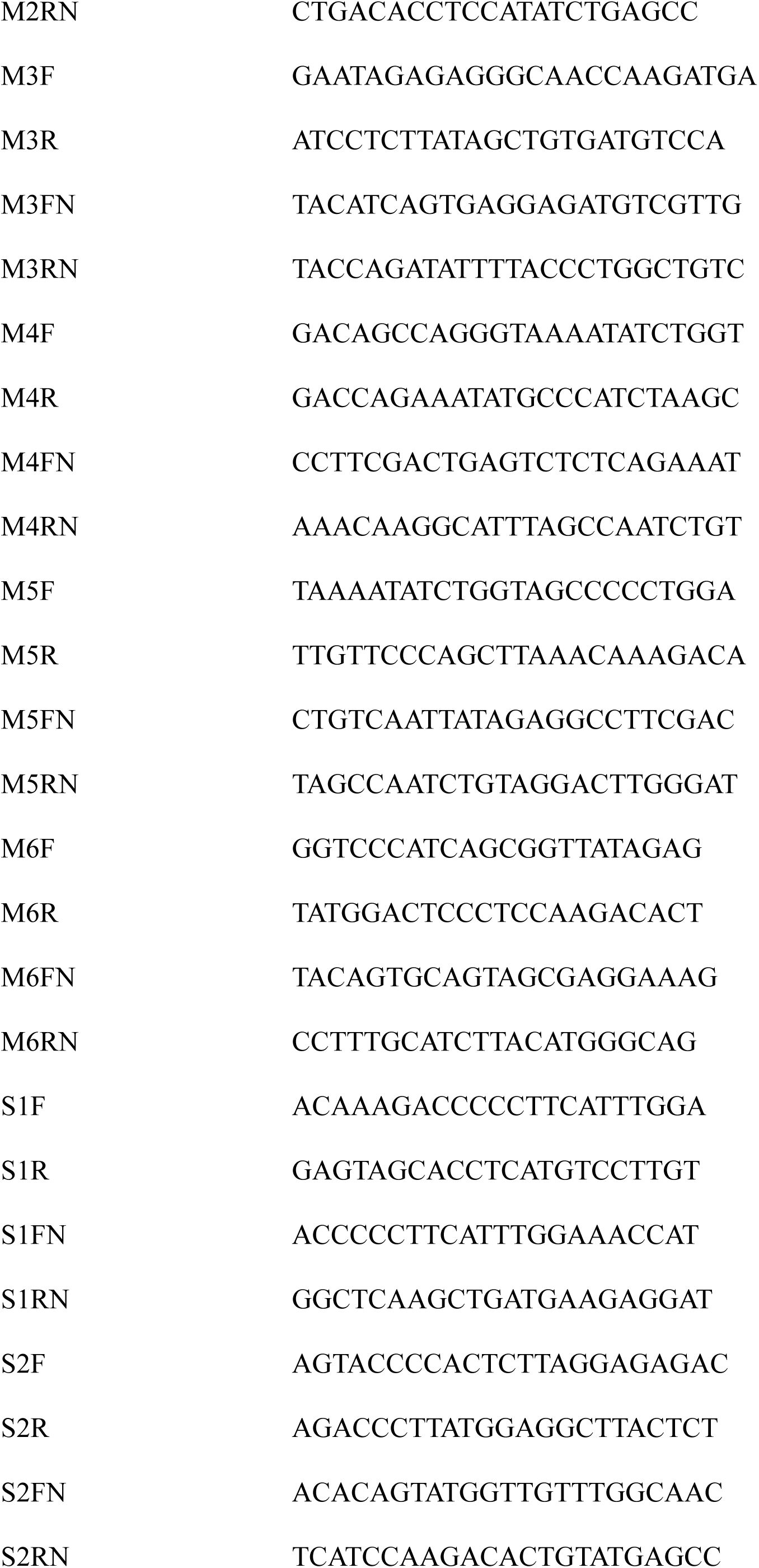
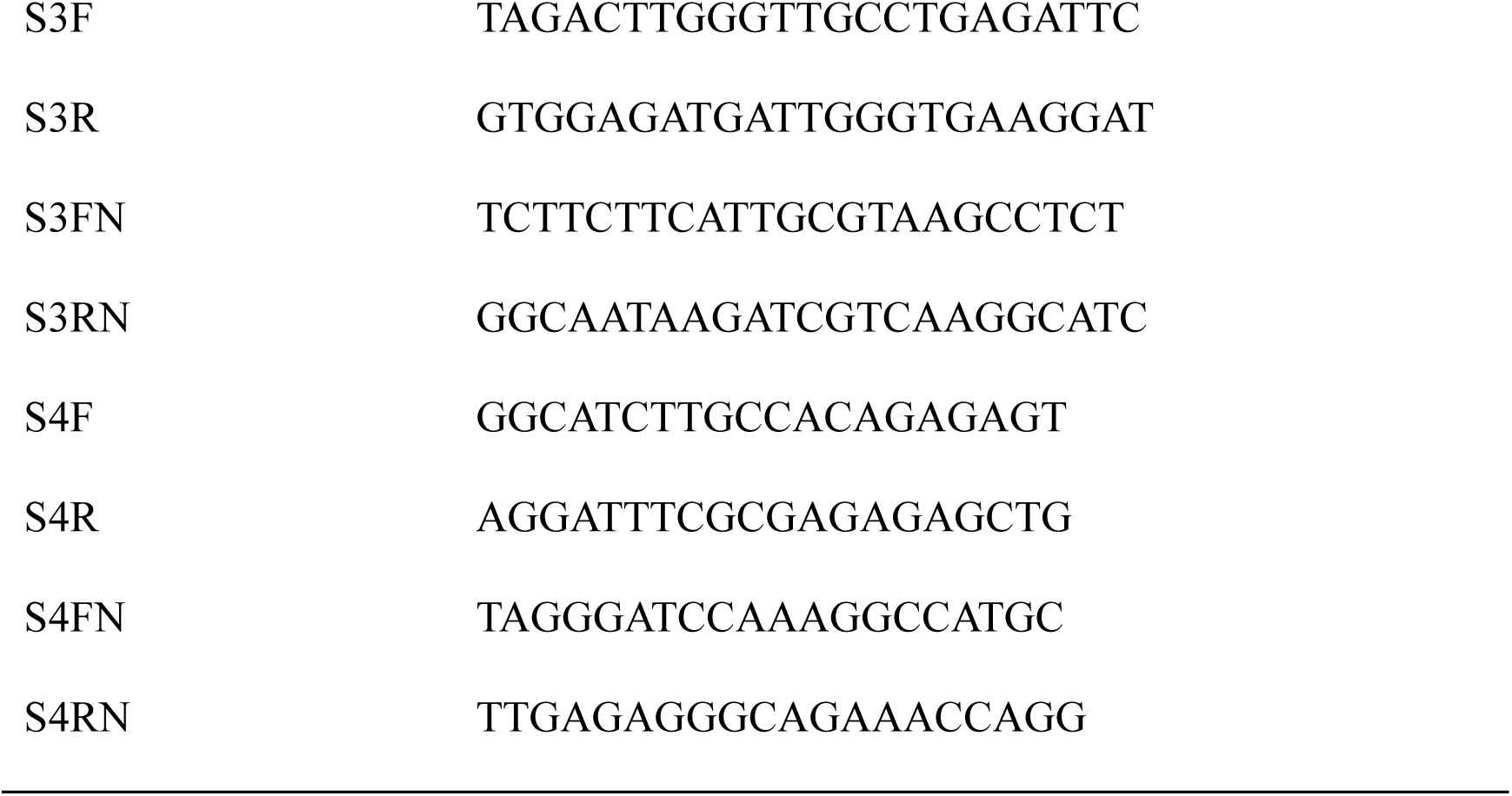
Nested-PCR Primers for SFTSV.

**Table S3.** GenBank accession numbers of SFTSV strains in the phylogenetic analysis.

| Strains | Segment L | Segment M |
| --- | --- | --- |
|  | Accession No. | Accession No. |
| AH12/China/2010 | HQ116417 | HQ141590 |
| HN6 | HQ141595 | HQ141596 |
| SDLZtick12/2010 | JQ684871 | JQ684872 |
| LN2 | HQ141607 | HQ141608 |
| HB155/China/2011 | JQ733564 | JQ733563 |
| HB29 | HM745930 | HM745931 |
| HB154/China/2011 | JQ733561 | JQ733560 |
| HB156/China/2011 | JQ733567 | JQ733566 |
| JS4 | HQ141604 | HQ141605 |
| SD4 | HM802202 | HM802203 |
| HN13 | HQ141598 | HQ141599 |
| HNXY206 | KC292329 | KC292302 |
| HNXY 191 | KC292349 | KC292323 |
| HNXY 278 | KC292348 | KC292322 |
| SPL097A_(Hyogo_2013) | AB983518 | AB985312 |
| SPL100A_(Hyogo_2013) | AB983519 | AB985313 |
| SPL057A_(Yamaguchi_2013) | AB983500 | AB985295 |
| SPL127A_(Miyazaki_2013) | AB985641 | AB985654 |
| SPL129A_(Miyazaki_2014) | AB983533 | AB985326 |
| SPL077A_(Ehime_2013) | AB983510 | AB985304 |
| SPL055A_(Yamaguchi_2013) | AB983499 | AB985294 |
| SPL075A_(Kochi_2013) | AB983509 | AB985303 |
| SPL078A_(Kumamoto_2013) | AB983511 | AB985305 |
| SPL062A_(Kumamoto_2013) | AB983502 | AB985297 |
| AHZ-2019-WXP | OP651054 | OP729245 |
| AHZ/China/2011 | JQ670929.1 | JQ670931.1 |
| 2022HNYGS8 | OP652095.1 | OP652067.1 |
| BJ001-2021 | OQ832099 | OQ832098.1 |
| Houjiazhuang Tick | C_AA059120.1 | C_AA059144.1 |
| Gaoying Patient | C_AA059131.1 | C_AA059145.1 |
| Beishiqu Tick | C_AA059142.1 | C_AA059146.1 |
| Beihai Tick | C_AA059153.1 | - |
| Wanwanshu Tick | C_AA059127.1 |  |
| Beiwu Tick | C_AA059154.1 | - |
| Jingshan Tick | C_AA059121.1 | C_AA059129.1 |
| Jingshan Hedgehog | - | C_AA059130.1 |
| Beihai Hedgehog | - | C_AA059132.1 |
| Zizhuyuan Hedgehog | - | C_AA059136.1 |
| Zizhuyuan Tick | - | C_AA059137.1 |
| Jishuitan Hedgehog | - | C_AA059138.1 |
| Wanliu Hedgehog | - | C_AA059140.1 |

