## Supplementary figures and images for "The first discovery of SFTSV in the Centre of Metropolitan Beijing, China"

### ciwei (1).tif

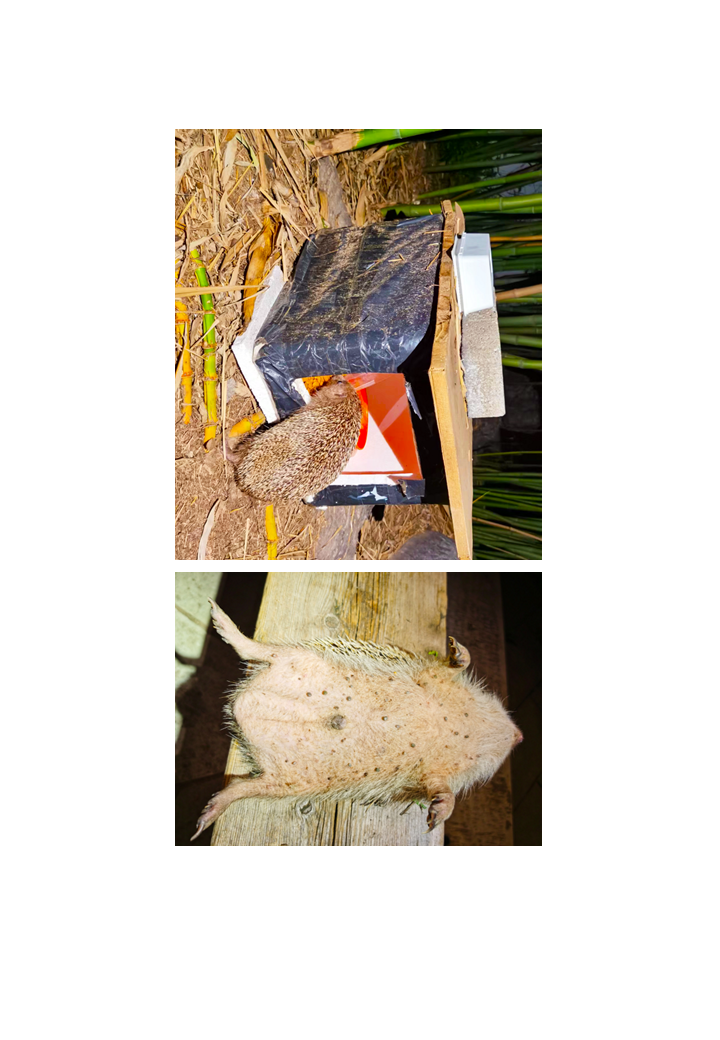

### figS1.tif

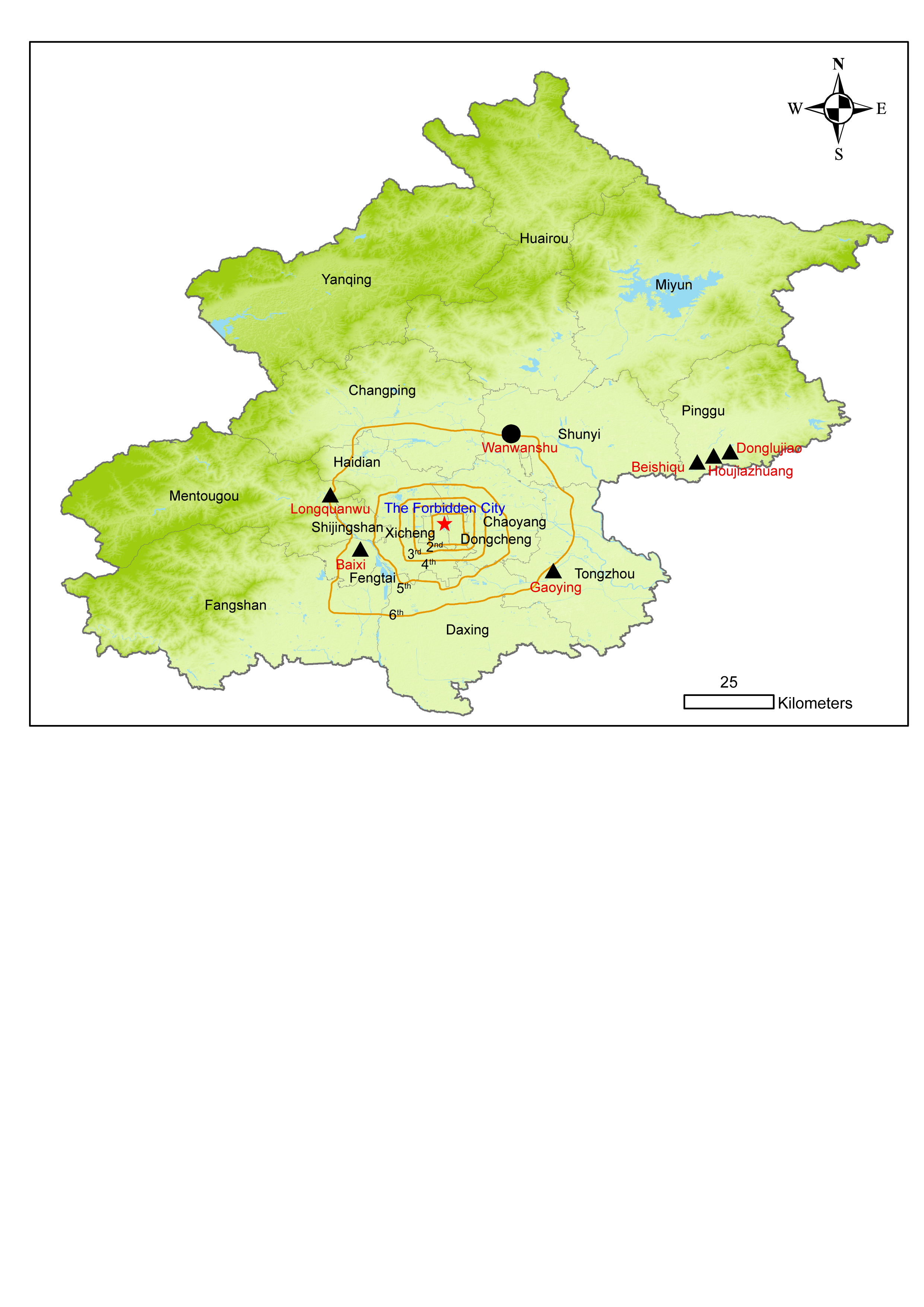
